# Mutualism mediates legume response to microbial climate legacies

**DOI:** 10.1101/2024.11.18.624153

**Authors:** Julia Anne Boyle, Bridget Murphy, Fangming Teng, Parsa Babaei Zadeh, Ingo Ensminger, John R Stinchcombe, Megan E Frederickson

## Abstract

Climate change is altering both soil microbial communities and the ecological context of plant-microbe interactions. Heat, drought, and their legacies can alter soil microbiomes and potential plant symbionts, but the direct consequences of these microbial changes on plant performance and plant investment in symbiosis remain underexplored. Predicting how soil microbes modulate plant resilience to heat and drought is critical to mitigating the negative effects of climate change on ecosystems and agriculture. In this study, we conducted growth chamber experiments to isolate the microbially-mediated indirect effects of heat and drought on plant performance and symbiosis. In the first experiment, focused on drought, we found that drought and drought-treated microbes, along with their interaction, significantly decreased the biomass of *Medicago lupulina* plants compared to well-watered microbiomes and conditions. In a second experiment, we then tested how the addition of a well-known microbial mutualist, *Sinorhizobium meliloti*, affected heat- and drought-treated microbiomes’ impact on *M. lupulina*. We found that drought-adapted microbiomes negatively impacted legume performance by increasing mortality and reducing branch number, but that adding rhizobia erased differences in plant response to climate treated soils. In contrast, heat-adapted microbiomes did not differ significantly from control microbiomes in their effects on a legume. Our results suggest microbial legacy effects, mutualist partners, and their interactions are important in mediating plant responses to drought, with some mutualists equalizing plant responses across microbial legacies.

## Introduction

How plant-microbe symbioses respond to climate change is hard to predict (Classen et al. 2015, Rudgers et al. 2020) because plants and microbes are each affected by climate directly and indirectly through their interactions (Figure 1) (Allison and Treseder 2008, Castro et al. 2010, Sheik et al. 2011, Franklin et al. 2016, Melillo et al. 2017, Zhou et al. 2020, Pec et al. 2021, Seaton et al. 2021). Plants and microbes cycle nutrients such as nitrogen and phosphorus, often while engaged in symbiosis, so if these relationships are altered by components of climate change like excessive heat and drought, ecosystem function may be altered (Kardol et al. 2010). In addition, because microbes can strongly affect plant fitness, outcomes of symbiosis are critical for predicting plant performance and evolution under climate change (Classen et al. 2015, Hawkes et al. 2020). Here, we experimentally tested the microbially-mediated indirect effects of heat and drought on plant performance and symbiosis.

**Figure 1.**
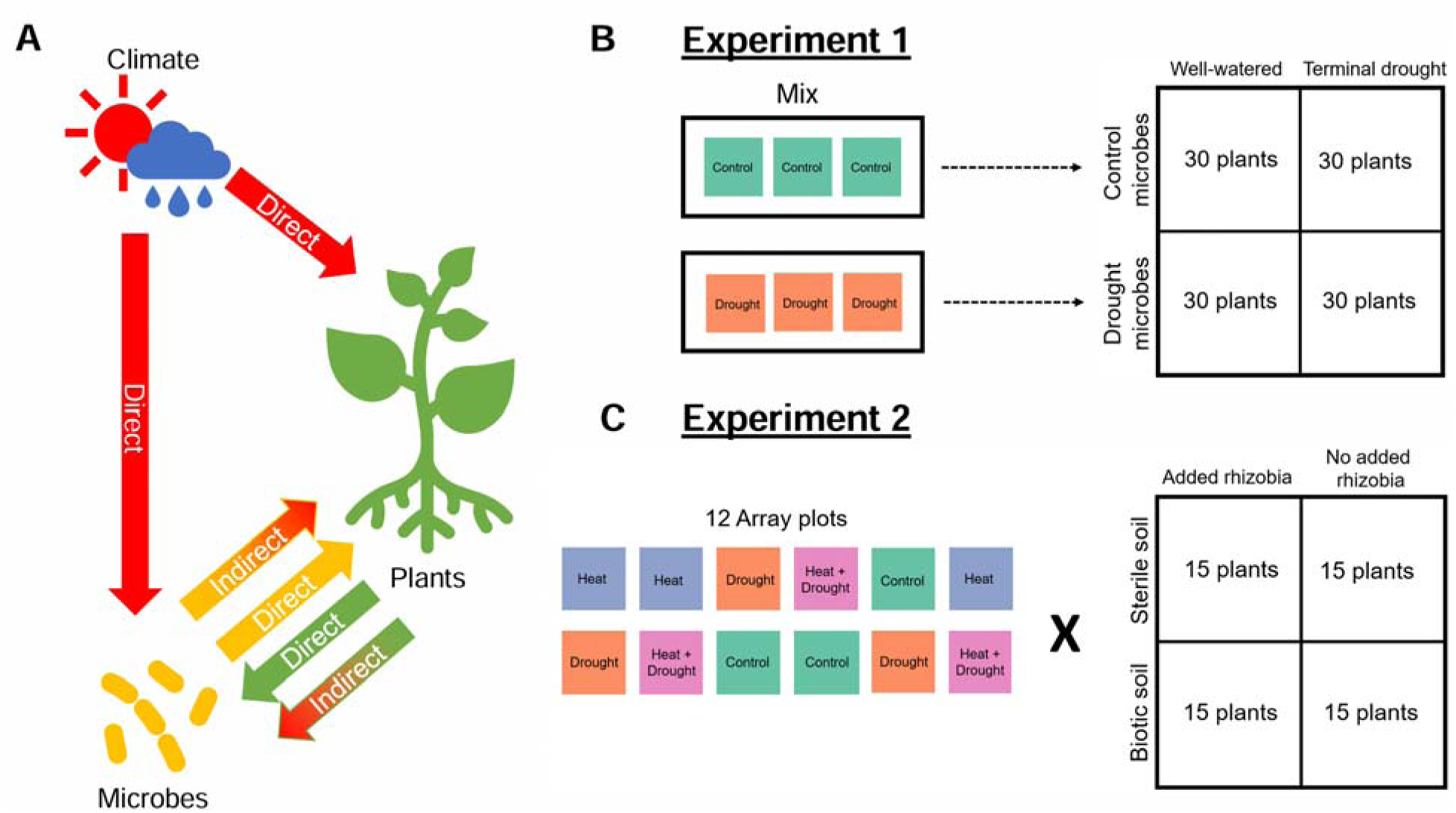
Conceptual motivation and treatment design for the two experiments. A: Plants and microbes experience the direct effects of climate (fully red arrows), as well as the indirect effects of climate mediated through plant-microbe interactions (partially red arrows). Here, we investigate the indirect effect of climate on plants, mediated through the microbiome (yellow to red arrow). B: We pooled soil from control and drought-treated field plots and added them to plants in well-watered or terminal drought conditions. C: For soil from each field plot, we factorially tested the effect of biotic vs. autoclaved soil and added rhizobia vs. no added rhizobia.

Soil microbiomes respond to heat and drought through changes in microbial diversity and community composition (Allison and Treseder 2008, Castro et al. 2010, Melillo et al. 2017, Pec et al. 2021), and the response of symbiotic microbes may not be in the same direction or of the same magnitude as in the broader microbial community (Boyle et al. 2024). Changes in a soil’s microbial community composition can alter rhizosphere microbiomes and the benefit that rhizosphere microbes provide to plants (Rubin et al. 2018, Lu et al. 2018, Fitzpatrick et al. 2019) with reproducible effects (Panke-Buisse et al. 2015), although elucidating the links between climate-driven changes in the soil and changes in the rhizosphere’s benefit is still needed. In addition to shifts in community composition, soil microbes may also respond to climate stressors via within-strain evolution or plasticity (Sachs et al. 2011, Simonsen 2021), and potentially push plant-microbe interactions along the mutualism-parasitism continuum (Davis et al. 2015, Shantz et al. 2016, Rudgers et al. 2020, González et al. 2021). Whether multiple climate stressors have compounding effects on soil microbiomes and plant-microbe symbioses is infrequently tested (Kivlin et al. 2013), but remains important given that multiple climatic axes are changing simultaneously (Thuiller 2007).

Determining the importance and scale of direct versus indirect effects of climate on plant-microbe symbioses is challenging if both direct and indirect effects are occurring at the same time. To address indirect effects only, studies can leverage the microbial legacy effects left behind by climate stressors (Averill et al. 2016, Martiny et al. 2017, de Nijs et al. 2019). Most studies examining the effects of climate-treated microbes suggest that they tend to improve plant performance in conditions that match those they evolved in (Lau and Lennon 2012, Allsup et al. 2023, Ricks and Yannarell 2023); these studies demonstrate that indirect effects of climate change on plants, mediated through microbial symbionts, may contribute to plant performance under changing climates. Yet shifts in the microbiome due to climate change have the potential to be either beneficial or detrimental to plants. On the one hand, microbiomes may facilitate a faster adaptive response than relying solely on host evolution, since microbiomes can rapidly change in composition (Copeland et al. 2015, Sepulveda and Moeller 2020) and this may offer new or more robust beneficial functions to the host (Kolodny and Schulenburg 2020). On the other hand, changes in the microbiome could also lead to an increase of mismatched or harmful symbiotic partners (Delgado-Baquerizo et al. 2020).

In a previous study, Boyle et al. (2024) factorially implemented heat and drought treatments on controlled field plots, and characterized changes in the communities of potentially symbiotic microbes found in the bulk soil. Known bacterial symbionts showed a strong response to climate treatments, including rhizobia, which form a specialized structure on legume roots called a nodule, where they fix nitrogen in exchange for fixed carbon (Beringer et al. 1979). Drought increased the diversity of both rhizobia and plant beneficial bacteria (PBB) and shifted phytopathogen community composition. The increased diversity of rhizobia and PBBs could mean that there are more potential strains for a plant to associate with, however given the high specificity of legume-rhizobium interactions (Chen et al. 2021), increased diversity could also make it harder for legumes to find compatible, high-quality partners (Burghardt and diCenzo 2023). Fungi also had strong community-wide responses to climate treatments, with mycorrhizal abundance decreasing in heat-treated plots and parasitic fungi increasing in drought-treated plots (Boyle et al. 2024). Mycorrhizae can increase plant stress tolerance (Quiroga et al. 2017; Kakouridis et al. 2022), provide additional nutrients (Koide 1991), and act as ‘fungal highways’ for bacteria to travel along to the plant root (Zhang et al. 2020). Thus, the previous results suggest heat and drought-induced variation in the soil microbiome could have a variety of consequences for plant performance.

The role of mutualism in buffering plants against the effects of climate change remains an open empirical question (Keeler et al. 2021, Duarte and Maherali 2022). If mutualistic symbionts stabilize plant fitness during times of fluctuating environment or microbiome quality (Mendes et al. 2011, Porter and Sachs 2020), this could affect how plants evolve and respond to climate change. Additionally, the presence of beneficial microbes in climate-treated soils may alter the magnitude of microbially-mediated indirect effects of climate change on plants, although to our knowledge, this has not been tested. How mutualisms buffer partners against climate change also has implications for the maintenance of mutualism (Kiers et al. 2010). For example, given stressful conditions, mutualists may lose one or more partners, switch to antagonism or new partners, or abandon mutualism altogether (Sachs and Simms 2006, Kiers et al. 2010). Whether heat or drought facilitates any of these outcomes may be detectable in clearly defined partnerships, such as the nutritional mutualism between legumes and rhizobia.

We experimentally tested the indirect effects of heat and drought on legumes mediated through changes in the soil microbiome, and whether the addition of rhizobia buffered legumes against these microbiome effects. Given the large impact of drought on the soil microbial community (Boyle et al. 2024), we first tested the effect of control and drought-treated microbes under both drought and well-watered conditions. We predicted that drought-treated microbiomes would improve legume tolerance of drought stress compared to control microbiomes, since microbes in the soil should be robust to drought, but that drought-treated microbes would be worse for plants under well-watered conditions. Then, in a second experiment, we added soil from the plots that experienced heat, drought, both, or neither to legumes in a growth chamber with and without added rhizobia, and quantified plant performance and plant investment in symbiosis. We predicted that adding climate-treated microbes to plants would decrease plant performance and weaken plant investment into symbiosis, but that adding rhizobia would buffer against these effects. Within this second experiment, we used 16S and ITS amplicon sequencing of rhizosphere and nodule microbes, in combination with previously published 16S and ITS data on the climate-treated bulk soil microbial communities used as inocula, to understand the impact of heat and drought legacy on plant microbiomes.

## Methods

### Study system

The temperature-free-air-controlled enhancement (T-FACE) experiment is in an old field at Koffler Scientific Reserve (KSR, www.ksr.utoronto.ca) in Ontario, Canada (44°01’48”N, 79°32’01”W). The experimental design, treatment effectiveness, and soil microbiome analysis are fully described in Boyle et al. (2024), but we give a brief overview here. Plots in the T-FACE experiment grew only seedlings of white spruce, *Picea glauca*, and were either heated, droughted, heated and droughted, or ambient (3 plots/treatment). Boyle et al. (2024) applied treatments during the growing seasons of 2020 and 2021, with rainout structures present for 8 months and heaters activated for 9 months. During active treatment, the mean soil temperature of heated plots was 3.7 or 3.6 hotter than un-heated plots in 2020 and 2021, respectively. In 2020, the mean soil volumetric water content (VWC) during droughts was 0.28 (m^3^/m^3^) in non-drought plots and 0.25 (m^3^/m^3^) in drought plots, and in 2021, mean VWC was 0.26 (m^3^/m^3^) in non-drought plots and 0.21 (m^3^/m^3^) in drought plots. We collected sifted bulk soil (passed through a 4.75 mm metal sieve) from each plot on June 5th, 2022, sterilizing tools between each collection. We kept soil bags stored slightly open in the dark at 4 until application to the experimental legumes.

We used seeds of a single genotype of *Medicago lupulina*, an annual legume that forms indeterminate root nodules with rhizobia in the genus *Sinorhizobium* (Batstone et al. 2020b). Seeds were collected from KSR and selfed for two generations in the University of Toronto greenhouses (Simonsen and Stinchcombe 2014). *Medicago lupulina* is naturalized at KSR near the experimental warming array, and its growing season peaks in June. The growth chamber experiments were not designed to mimic ecological interactions in nature, but rather to test the general hypothesis that soil microbial changes, including known changes in rhizobia in response to heat and drought (Boyle et al. 2024), can by themselves affect plant performance and symbiosis. We chose the legume *M. lupulina* for experimental tractability and because it partakes in multiple microbial symbioses, including symbiosis with rhizobia and arbuscular mycorrhizal fungi.

### Experiment 1: Effect of drought-treated microbes on legume performance in drought

#### Experimental design and data collection

We tested whether drought-treated microbes affected legumes in both dry and well-watered soil conditions, because past work showed that drought was particularly important for shifting microbial community composition (Boyle et al. 2024). We implemented a 2×2 factorial design with plants receiving biotic (e.g. live) drought soil or biotic control soil, and experiencing a terminal drought or well-watered conditions, with 30 replicates per treatment, for a total of 120 plants. We pooled soil within treatment types to generate inocula. We surface-sterilized seeds with bleach and ethanol then stratified them for two weeks at 4 °C. We planted them into Magenta boxes containing autoclaved Profile® fine-grained turface and a nylon rope wick, spatially blocked by light canopy in the growth chamber. The experiment took place in a growth chamber room set to cycle between 22 °C day/20 °C night with 16 hours of light.

Upon planting, we applied 10 ml of soil to each plant. We added soil directly to maintain the integrity of fungal structures and the diversity of microbes. Additionally, we applied 1 ml of low-nitrogen fertilizer (Supplemental material) to each plant, and we reapplied 1 ml of fertilizer every 4 weeks. All pots were covered with plastic wrap to maintain moisture, and when covered plants grew tall enough, we made a hole in the cover to accommodate their growth (Supplemental Figure 1). Four weeks post-planting, we uncovered the terminal drought plants to allow more drying and kept the well-watered plants covered. We filled the bottom compartments of well-watered plants with 300 ml of autoclaved dH_2_O. We began the gradual terminal drought treatment with 2 weeks of well-watered conditions, followed by 2 weeks of ⅔ volume water, then 3 weeks of ⅓ volume water, and finally no further water. We refilled all plant water compartments with the appropriate volumes whenever the well-watered plants used all their water. Ten weeks post-planting, just over 80% of terminal drought plants were dead and we ended the experiment. We counted branch number approximately every 10 days, and measured the nodule number and dry weight of above- and below-ground biomass of each plant, including dead plants, at the end of the experiment.

#### Statistical analysis

We fit linear models with log_10_-transformed above- and below-ground biomasses as response variables and the main and interactive effects of soil origin and experimental water condition as fixed effects and spatial block as a random effect. Final branch number was tested in a generalized linear model with a Poisson distribution. We included data from both live and measurable dead plants in the models. Only a single plant made nodules, and thus we did not fit a statistical model to nodule number.

### Experiment 2: Soil climate legacy effects with and without an additional mutualist

#### Experimental design

In a separate experiment, we next tested whether heat- and drought-treated microbes affected legumes and rhizobia, under well-watered conditions only. We factorially tested the four climate treatments, each with 3 replicate plots from the warming array, and with sterilized or fresh soil. The same collection of soil was used in Experiment 1 and 2. We used autoclaved sterilized soil (Supplemental Material) to test for abiotic differences in soil from different field plots and climate treatments. The field soil resulted in poor nodulation in Experiment 1, so, in Experiment 2, we also inoculated half of the plants with *Sinorhizobium meliloti* 1021-71 tagged with green fluorescent protein (GFP) (courtesy of Daniel Gage, Gage et al. 1996) to determine whether additional rhizobia buffer legumes against the negative effects of drought-treated microbiomes. In total, we had 48 treatments replicated 15 times for a total of 720 plants (Figure 1). We also included an additional 15 sterile control plants with no additional soil.

We prepared the same genotype of seeds as before and blocked the plants by light canopy under the same growth chamber conditions as previously described. We placed cone-tainers with a wicking nylon rope and 130 ml of autoclaved Profile® fine-grained turface into 50 ml falcon tubes with autoclaved distilled water, and refilled as needed. We added 1 ml of low-nitrogen fertilizer upon planting, two weeks after planting, and every 3 weeks subsequently. We applied biotic and sterilized soil treatments five days after planting, and applied *S. meliloti* 1021-71 or sham inoculations six days after planting, as described in the Supplemental Material. Only two sterile plants in the experiment made nodules, indicating minimal rhizobial contamination. Autoclaved soil, however, was very susceptible to ambient fungal colonization, and 154 sterilized soil replicates showed mold growth. We removed all plants that showed signs of contamination from further analysis.

#### Data collection

Over almost 12 weeks, we surveyed branch number and mortality. At the end of the experiment, we randomly sampled five live plants per treatment receiving biotic soil to sequence the microbial communities in their rhizosphere and nodules (n = 120). We did not use fluorescence to determine rhizobia strain identity in nodules because of the possibility of bacterial autofluorescence (Yang et al. 2012) and because we sequenced nodule bacteria. As described in the Supplemental Material, we extracted DNA from the rhizosphere and nodule microbes, then sequenced the fungal ITS region and bacterial 16S V4 region using Illumina MiSeq paired end sequencing. For all plants, we counted the number of nodules then dried and weighed above and below ground biomass.

#### Statistical analysis: Plant phenotypic responses

We initially tried to fit a full four-way ANOVA (heat * drought * sterility * rhizobia) with random effects of array plot and block to our data, but found that these models did not converge (or had error warnings) for some of our response variables, precluding the use of a single model structure for all the phenotypes we measured. As such, we tested the effects of added rhizobia and soil sterility separately from the effects of climate (heat and drought treatment). In the analyses described below, our inferences about rhizobia mediating plant responses to climate legacies come from testing for significant differences due to climate in biotic soils, with or without additional rhizobia (Figure 1C); because the only difference between these treatments is the presence or absence of rhizobia addition, we attribute the differences to rhizobia.

First, to test how added rhizobia and soil sterility affected plant outcomes, we used a generalized linear mixed-effects model with sterility treatment and rhizobia treatment as interacting fixed effects, and block and array plot as random effects. As an exception, to test nodule data we used a zero-inflated Poisson model in the GLMMadaptive package (Rizopoulos 2023) with heat, drought, and block as fixed effects, and array plot as a random effect; we removed the “sterile no added rhizobia” treatment group from this analysis because its values were only zero. The zero-inflated model included both a model of the zeros (nodule presence or absence), and a model of the non-zeros (nodule number).

Then, we tested for effects of soil climate origin within each of the four broad treatment categories using generalized linear models with heat and drought as interacting fixed effects, and with block and array plots as random effects, with exceptions and modifications described below. To test how climate-treated microbes affected plant performance, we used repeated branch counts as the response variable, with an additional fixed effect of days since germination and random effect of unique plant ID. We tested branch count in the biotic no added rhizobia and biotic added rhizobia treatments separately. In the biotic soil with no added rhizobia treatments, to test nodule data, we used a zero-inflated Poisson model with heat, drought, and block as fixed effects and array plot as a random effect.

We normalized response variables as needed or used generalized linear models. We log_10_-transformed above- and below-ground biomasses and nodule number, and log_10_(y+1)-transformed leaf counts to use a Gaussian family. We tested mortality and nodule presence with a binomial family. We performed analysis in R v4.2.0 (R Core Team 2022), with tidyverse (Wickham et al. 2019), cowplot (Wilke 2017), lme4 (Bates et al. 2015), and lmerTest (Kuznetsova et al. 2017). We used type 3 ANOVAs from the car (Fox and Weisberg 2011) package to calculate *Wald X^2^* values and significance for all linear and generalized linear models. We excluded plants that did not successfully germinate as well as plants with suspected contamination from the analysis.

#### Statistical analysis: Plant microbiomes

We tested the effects of climate legacy and added rhizobia on the rhizosphere and nodule microbial communities, and linked microbes to plant performance. We analyzed nodule, bacterial, and fungal rhizosphere datasets separately, using the phyloseq (McMurdie and Holmes 2013), and microbiome (Lahti and Shetty 2019) packages. We excluded any potential root endophytes from our nodule samples by subsetting reads to include only *Sinorhizobium* ASVs. We used generalized linear models with heat and drought as interacting fixed effects, the fixed effect of added rhizobia, and random effects of block and array plot; when we tested nodule data we added total nodule number as a fixed effect. In the rhizosphere data, we tested observed richness, Shannon diversity, and evenness using Gaussian distributions. To test alpha diversity in nodules, we used a Poisson distribution for observed richness, a Gaussian distribution for evenness, and a binomial distribution for Shannon diversity (due to the frequency of a single ASV being observed, we categorized Shannon diversity as 0 or > 0). Next, we tested how climate legacy affected microbial composition using adonis2 (vegan package, (Oksanen et al. 2020) permutational analysis of variance models (PERMANOVA; Bray-Curtis method, 9999 permutations) with heat and drought as interacting fixed effects, added rhizobia as another fixed effect, and permutations done within blocks. To test nodule composition, we added total nodule number as a fixed effect.

Finally, to identify microbes correlated to plant performance, we subset the top 20% best performing (n = 23) and bottom 20% worst performing (n = 23) plants from the sequenced plants after rarefaction in both the bacterial and fungal datasets. Then, using the microviz package (Barnett et al. 2021), we used the tax_fix function, center log ratio transformed abundances of the rarefied rhizosphere reads, and used a principal components analysis (PCA) to plot points and identify the top 10 genera with the largest effect sizes. We tested whether the best and worst performing plants had significantly different rhizospheres using a PERMANOVA (Aitchison method, 9999 permutations accounting for block). To match our PCA, we aggregated taxa abundance to genus, then used performance category and added rhizobia as fixed predictors, and heat and drought as interacting predictors.

## Results

### Drought-treated microbes reduced legume performance

In both experiments, drought-treated microbes harmed plants. In Experiment 1, plants grew largest under well-watered conditions, but only when inoculated with soil microbes from the control, not the droughted, field plots. There were significant effects of both current water condition and the historical soil water condition on aboveground biomass (*Wald X^2^* > 18.5, *p* < 0.001 for all effects; Supplemental Table 1, Figure 2A), with a significant interaction (*Wald X^2^* = 5.98, *p* < 0.05; Supplemental Table 1, Figure 2A). When plants received ample water, drought-treated microbes reduced plant aboveground biomass by 51.3% compared to control microbes, but in terminal drought conditions, drought-treated microbes reduced plant aboveground biomass by only 9.18% compared to control microbes. Belowground biomass also significantly decreased with drought-treated microbiomes (*Wald X^2^* = 18.0, *p* < 0.001; Supplemental Table 1, Figure 2B), and there was no main effect of the terminal drought treatment (Supplemental Table 1, Figure 2B), although there was a current conditions x historical conditions interaction (*Wald X^2^* = 7.44, *p* < 0.01; Supplemental Table 1, Figure 2B). The interaction term appears to be driven by the different responses to well-watered conditions between plants given control or drought-treated soil, when they had the same response in terminal drought (Figure 2B). The belowground biomass of plants with control soil decreased by 40.7% in terminal drought, compared to well-watered conditions. In contrast, plants with drought-treated microbes had 134% higher belowground biomass in terminal drought conditions, compared to well-water conditions.

**Figure 2.**
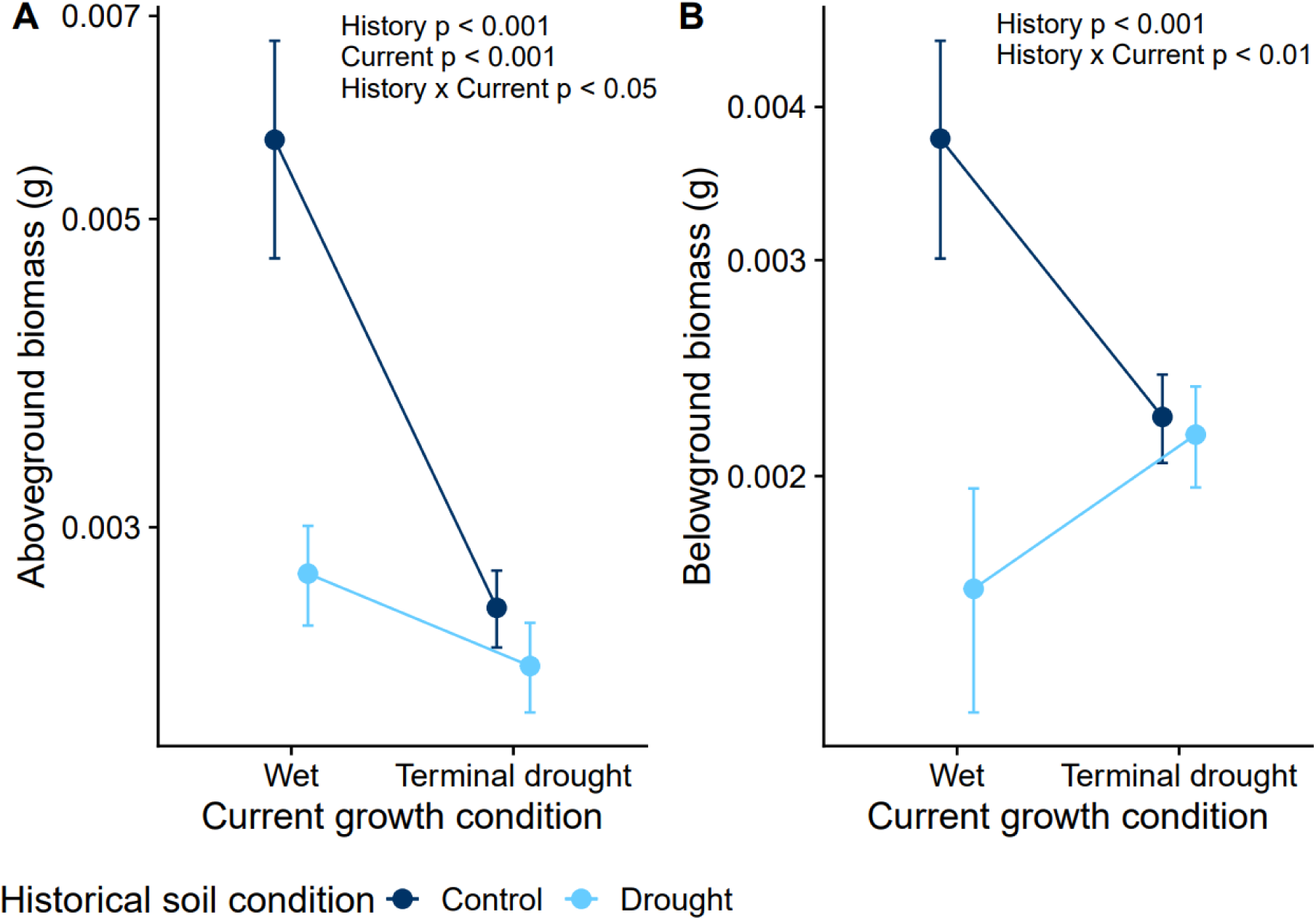
Climate legacy effects in soil microbiomes have context dependent effects on plants. Current and historical conditions, along with their interaction, significantly affected plant aboveground biomass (A) and belowground biomass (B) in the first experiment. Points are means with ±1 standard error. The y-axes are on a log_10_ scale.

Similarly, drought-treated microbes and terminal drought significantly decreased branch number (*Wald X^2^* > 4.64, *p* < 0.05 for all effects; Supplemental Table 1, Supplemental Figure 2), but did not significantly interact (Supplemental Table 1). Terminal drought also significantly increased mortality (*Wald X^2^* = 9.58, *p* < 0.01; Supplemental Table 1), but there were no mortality differences between microbe treatments (Supplemental Table 1). Only one plant made nodules, and it received control soil under well-watered conditions.

In our second experiment, conducted under well-watered conditions only, we found that *M. lupulina* performance was unaffected by heat-treated vs. control microbes, but again drought-treated microbes reduced plant performance. Plants with biotic, drought-treated microbes with no added rhizobia had a significantly reduced branch count compared to plants given biotic control microbes with no added rhizobia (*Wald X^2^* = 15.4, *p* < 0.001; Supplemental Table 2, Figure 3A). There was a significant interaction between drought and heat legacies on branch count (*Wald X^2^* = 5.37, *p* < 0.05; Supplemental Table 2, Figure 3A), where heat counteracted the negative effect of drought. In biotic soil with no added rhizobia, drought-treated microbes significantly increased mortality from 2.2% when inoculated with control microbes to 18% when inoculated with drought-treated microbes (*Wald X^2^*> 4.33, *p* < 0.05; Supplemental Table 2, Figure 3B).

**Figure 3.**
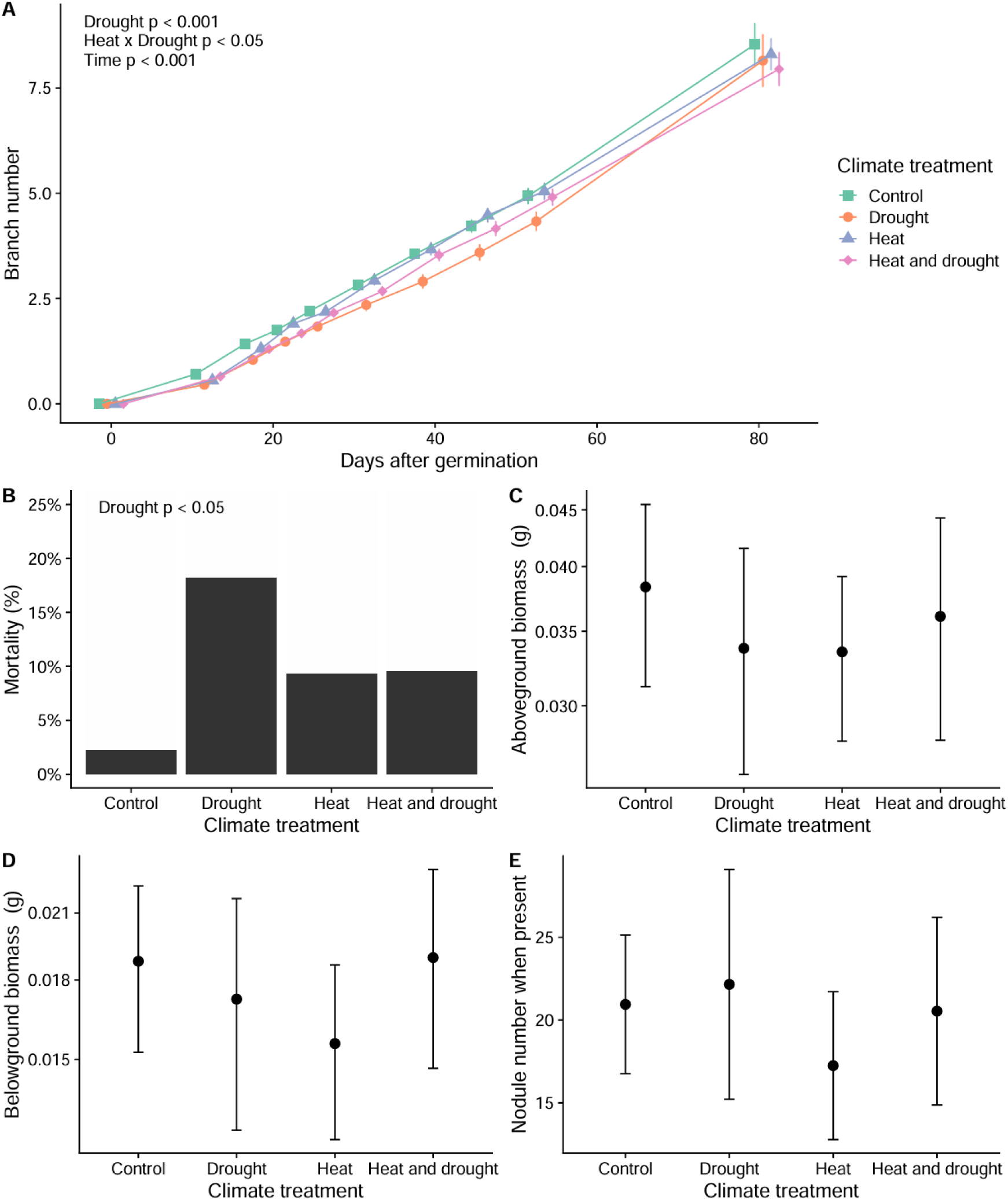
Drought-treated microbes decreased *Medicago lupulina* performance in the biotic, no added rhizobia treatment. In the second experiment, drought-treated microbes significantly decreased branch number when plants were young (A) and increased mortality (B), but by the end of the experiment, surviving plants had similar biomasses and nodule number across climate-treated soils (C, D, E). Points represent means ± 1 standard error. The y-axes in C and D are on a log_10_ scale.

However, by the end of Experiment 2, the above- and below-ground biomasses of surviving plants did not differ significantly among microbial treatments (Supplemental Table 2, Figure 3CD). While mean nodule number (including plants that made no nodules) decreased by 44% in plants inoculated with heat-treated field soil compared to control soil, the reductions in nodule presence and number was not statistically significant (Supplemental Table 2, Figure 3E). Differences in plant performance were not due to abiotic soil differences, because there were no significant differences in plant performance due to climate origin in the autoclaved, no added rhizobia soil treatments (Supplemental Table 3).

### Rhizobia reduced differences due to climatic legacy effects

Adding rhizobia to the biotic soil erased the significant plant performance differences that arose from inoculation with biotic drought-treated soils. In this treatment, mortality was 12-16% across climate treatments, with no significant differences due to climate legacy (Supplemental Table 4). Furthermore, there were no significant differences in branch counts, nodule number, nodule presence, or plant biomass among climate treatments in biotic soil with added rhizobia (Supplemental Table 4). There also were no differences due to microbial climate in the added rhizobia, sterilized soil treatment (Supplemental Table 5).

### Biotic soil improved plant performance more than adding rhizobia alone

When we tested the effects of soil sterility and added rhizobia, we found that plants performed best in unaltered field soil. Above- and below-ground biomasses significantly increased with added rhizobia inoculation (*Wald X^2^* > 5.23, *p* < 0.05 for all effects; Supplemental Table 6, Figure 4AB), but with a significant interaction between rhizobia and sterility condition (*Wald X^2^* > 6.64, *p* < 0.01 for all effects; Supplemental Table 6, Figure 4AB). Unsurprisingly, adding rhizobia increased plant biomass in sterilized soil, but adding rhizobia decreased plant biomass when plants were inoculated with biotic field soils. The mean aboveground biomass of plants in the biotic soil with the natural soil rhizobia community was 23% higher compared to plants in biotic soil with added rhizobia. Mortality was not affected by soil sterility or added rhizobia (Supplemental Table 6, Figure 4C). Nodule presence and number were significantly affected by soil sterility, added rhizobia, and spatial block (|*z-value*| > 2.04, *p* < 0.05 for all effects; Supplemental Table 6, Figure 4DE). Nodule presence decreased with no added rhizobia and no access to the natural rhizobia community. More nodules were made per plant with the unmodified field rhizobia, and plants made more nodules in the added rhizobia treatment only in the sterilized soil, when no other rhizobia were available.

**Figure 4.**
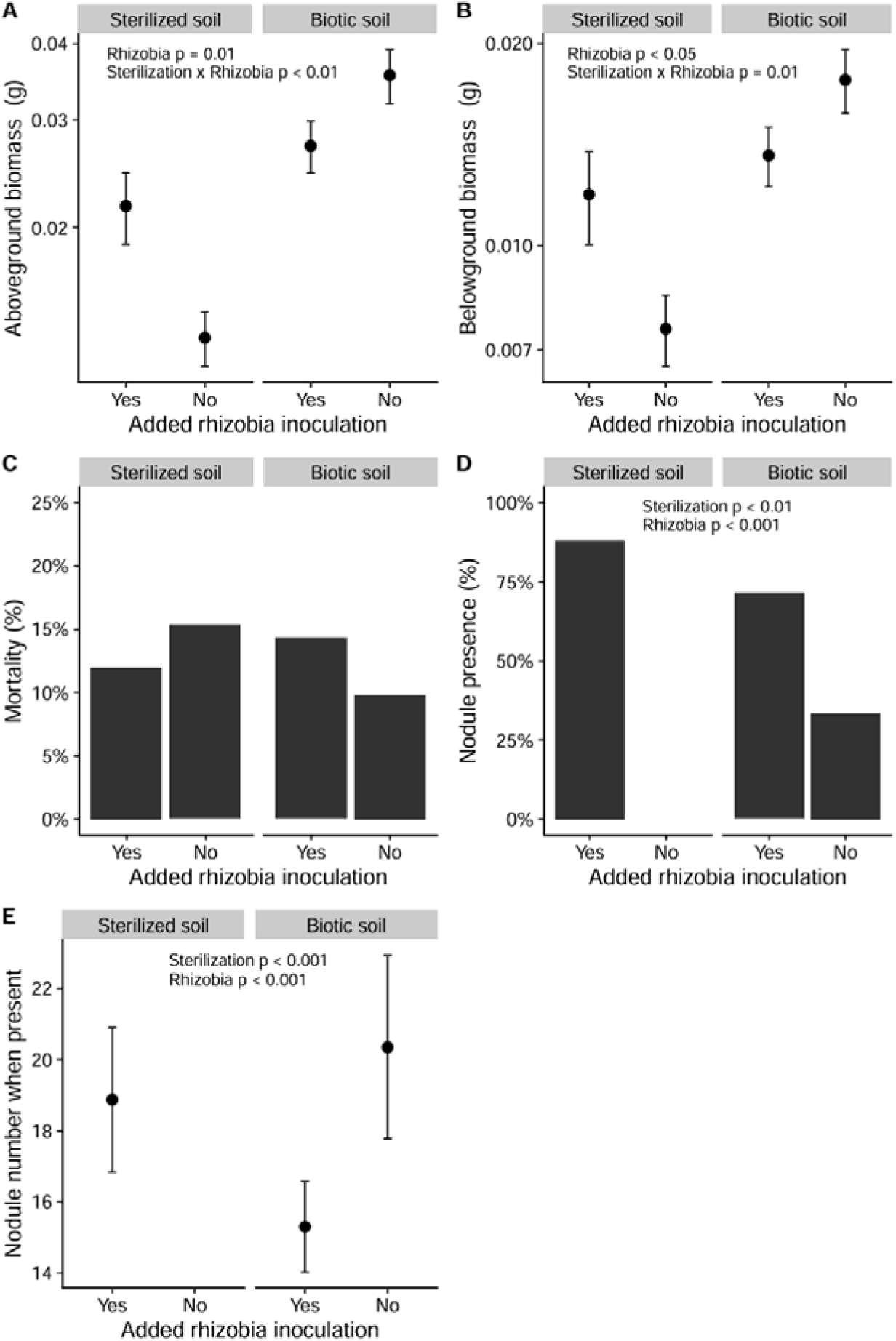
Plants performed best in the biotic soil with no added rhizobia treatments. In the second experiment, plants given unaltered field soil had the highest biomasses, while plants that received sterile soil with no additional rhizobia had the lowest biomass (A, B). The effect of adding rhizobia on biomass was dependent on the presence of other microbes, causing significant interaction terms (A, B). Mortality was not significantly different due to sterilized soil or added rhizobia (C). Nodule presence was highest when added rhizobia were the only microbes available, and second highest with added rhizobia and field microbes (D), but when plants did make nodules, they made the most with biotic unaltered field soil (E). Points represent means ± 1 standard error. The y-axes in A and B are on a log_10_ scale.

### Root microbiomes correlated to performance and changed by adding rhizobia

After rarefaction, bacterial rhizospheres contained 7,907 ASVs and had a high abundance of rhizobial taxa such as *Rhizobium* and *Bradyrhizobium*, but also high amounts of *Arthrobacter*, *Streptomyces*, and unknown taxa (Supplemental Figure 3). After rarefaction, fungal rhizospheres contained 992 ASVs, and were mainly composed of *Trichoderma* and *Fusarium*, with high occurrences of *Aspergillus*, *Neocosmospora*, and *Penicillium* (Supplemental Figure 4). Bacterial rhizosphere composition was not significantly altered by climate legacy (Supplemental Table 7) but was significantly altered by the addition of *S. meliloti* 1021-71 (*F*_1,108_ = 1.35, *p* < 0.05; Supplemental Table 7). Only heat-treated soils led to significantly higher observed bacterial richness in the rhizosphere compared to control soil (*Wald X^2^*= 4.08, *p* < 0.05; Supplemental Table 7), but Shannon diversity and evenness were not significantly different among treatments (Supplemental Table 7). Fungal rhizosphere community composition and diversity were not significantly affected by adding rhizobia or by the legacies of climate (Supplemental Table 8).

The best and worst performing plants had significantly different bacterial rhizospheres (*F*_1,39_ = 4.26, *p* < 0.001; Figure 5, Supplemental Table 7), but not fungal rhizospheres (Supplemental Table 8; Supplemental Figure 5). Of the 10 microbes with the largest effect, all but one were known plant beneficial bacteria (Li et al. 2023). The diazotrophs *Cellulomonas*, *Cryocola*, and *Sinorhizobium* (Hrynkiewicz et al. 2019) were positively correlated with plant performance (Figure 5). *Bacillus* was also positively correlated with plant performance (Figure 5) and can induce systemic resistance in plants, reducing the incidence and severity of phytopathogens (Kloepper et al. 2004). *Emticicia*, *Niastella*, *Variovorax*, and *Bdellovibrio,* as well as the families *Verrumicrobiaceae* and *Chitinophagaceae*, correlated with decreased plant performance (Figure 5).

**Figure 5.**
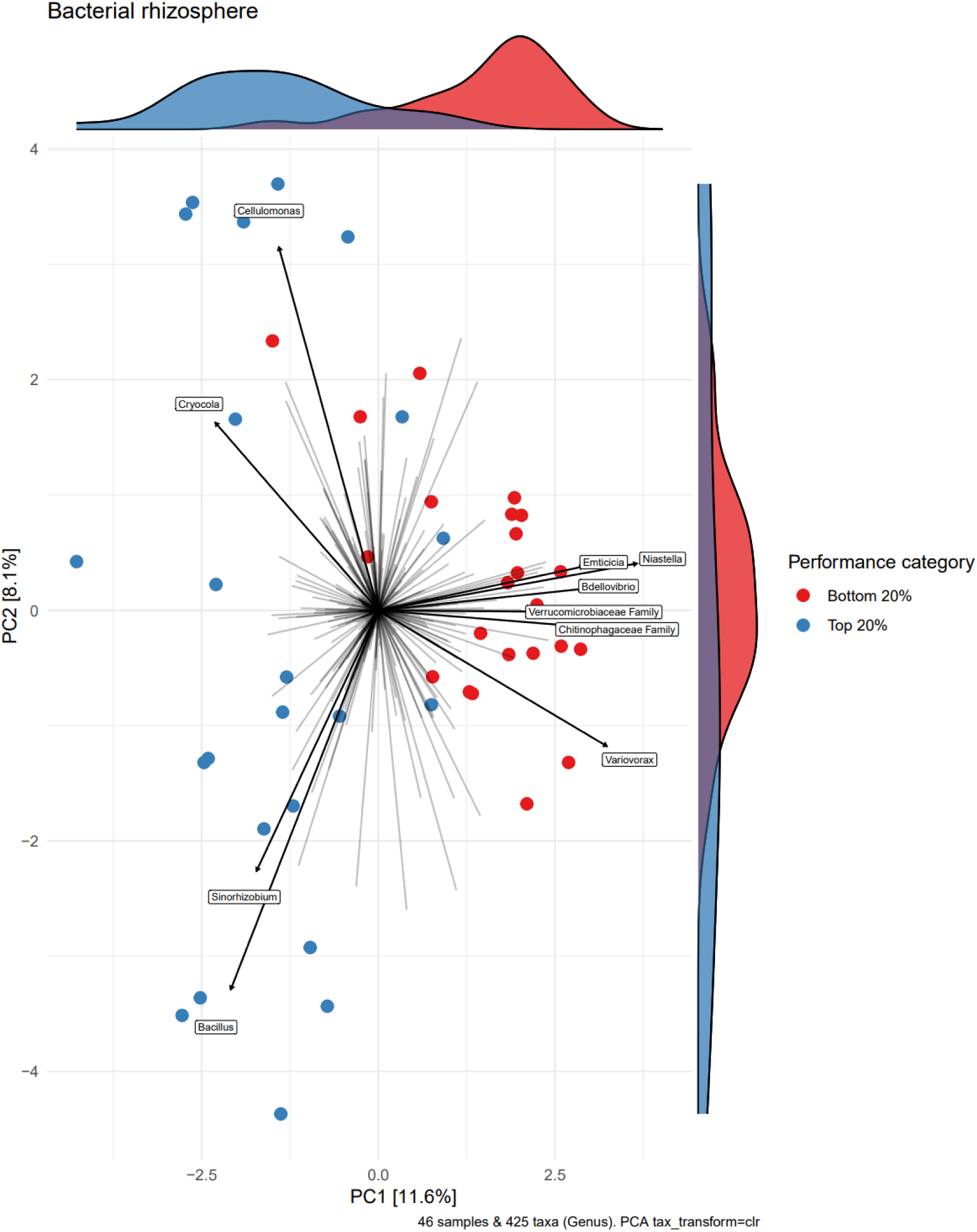
Microbes were correlated to performance differences in the bacterial rhizosphere. Here a PCA shows the center log ratio transformed bacterial rhizosphere composition of the top 20% best performing and bottom 20% worst performing plants in the second experiment. Performance was quantified by aboveground biomass (g).

Plants were very good at isolating *Sinorhizobium* from the soil (Supplemental Figure 3), and sequenced nodules were predominantly composed of 13 ASVs in the genus *Sinorhizobium*. *Sinorhizobium* ASVs were very similar to one another and to the ASV of *S. meliloti* 1021-71; we observed three different base pair substitutions, with other ASV diversity coming from 1-3 fewer or additional base pairs at the beginning and end of the sequence. The composition of those ASVs was not significantly affected by climate legacy, added rhizobia, or total nodule number (Supplemental Table 9). Increasing nodule number significantly increased the Shannon diversity of *Sinorhizobium* strains (*Wald X^2^*= 4.89, *p* < 0.05; Supplemental Table 9), but no other metrics of alpha diversity (Supplemental Table 9).

## Discussion

Climate events leave lasting effects on soil microbiomes, generating indirect microbially-mediated effects on plant performance and symbiosis. We found that drought-treated microbes negatively impacted legume performance in two experiments. In the first, drought-treated microbes significantly reduced plant biomass, and microbe history significantly interacted with experimental water availability to affect plant performance. In the second, drought-treated microbes increased mortality rates. When we added rhizobia, the differences in plant performance due to climate legacy became nonsignificant. Symbiosis outcomes were altered in other ways, with added rhizobia altering the bacterial rhizosphere composition and heat-legacy increasing the diversity of bacteria in the rhizosphere of live plants. Here we discuss the consequences of plant performance being shaped by these microbial indirect effects during climate change, how mutualism quality may remain stable in the face of climate change, and how mutualisms could potentially provide a stabilizing effect on plant performance across climate-driven differences in the microbiome.

### Negative effects of drought-treated microbes and their ecological implications

Plants respond to climate change through both direct and indirect pathways (Porter et al. 2020, Trivedi et al. 2022), and the relative importance and impact of these effects is necessary to predict plant performance and adaptation. Strikingly, the strength of microbially-mediated indirect effects of drought on plant performance was equal in magnitude to the direct effects of imposed drought (Figure 2). It is concerning that drought-treated microbiomes led to a worse plant performance because droughts are becoming more frequent and intense (Thuiller 2007, Allan and Soden 2008), and our results suggest that we cannot rely on soil microbiomes to contribute positively to plant resistance and resilience to climate change. Further, strong microbially-mediated indirect effects of climate change may impose additional ecological pressures on plants that must already respond to the direct effects of climate change. Previous studies found that drought microbiomes can buffer plants against drought stress (Lau and Lennon 2012, Allsup et al. 2023, Ricks and Yannarell 2023), but our results and recent results from Swift *et al*. (2024) indicate that this effect is not universal and that repeated drought events do not always select for microbes that improve plant performance relative to control microbes.

There are several possible mechanisms by which the drought-treated soil microbiomes may have decreased plant performance. The higher risk to plants could have been due to a lack of compatible mutualists, as the higher rhizobia diversity in drought (Boyle et al. 2024) may have increased competitive interference amongst rhizobia (Burghardt and diCenzo 2023, Rahman et al. 2023), lengthening the time needed for plants to locate compatible partners, and ultimately increasing risk of death. Given that plant performance was strongly positively correlated with *Sinorhizobium* abundance in the rhizosphere (Figure 5), and adding rhizobia erased climate-driven differences in plant performance, access to compatible rhizobia is clearly important for the plant. While the rhizobia diversity mechanism is formally plausible, we failed to detect other clear signals of it, such as corresponding differences in nodule number and diversity between surviving plants that received different climate-treated soils; as such, we think it is less likely.

Alternatively, our observed patterns of performance would occur if the prevalence of a detrimental microbe was higher in drought-treated soil. Indeed, the higher mortality may have been because of the 36% increase in relative abundance of fungal parasites and the altered bacterial phytopathogen composition in the drought inoculation soil (Boyle et al. 2024). Included amongst the plant fungal parasites in the inoculation soils are *Verticillium* and *Magnaporthiopsis*, which can cause wilt in legumes and non-legumes (Klosterman et al. 2009, Dor and Degani 2019), and *Paraphoma*, which can cause root rot in alfalfa and other plants using phytotoxins (Dang and Li 2022). Global warming and drought are projected to increase the proportion of soil-borne pathogens (Preece et al. 2019, Delgado-Baquerizo et al. 2020), and how potential pathogens interact with the rest of the plant microbiome under drought and heat stress is not well understood (Trivedi et al. 2022). Our results novelly suggest drought increases plants’ risk of encountering combinations of deadly microbes, but as we discuss more below, that the harm could be diluted if microbial mutualists are present at high enough frequency. Additionally, our sequencing results suggest that the microbes in the drought-treated soil and rhizospheres were less metabolically active and provided fewer nutrients to plants. The bacterial genus *Verrucomicrobium* was significantly enriched in the drought-treated inoculation soil (Boyle et al. 2024), and the *Verrucomicrobiaceae* family abundance in the rhizosphere was negatively correlated with plant performance in Experiment 2 (Figure 5). Possibly due to its slow growth, the phylum *Verrucomicrobia* is linked to reduced soil enzyme activity and loss of soil nutrients like nitrogen (Xie et al. 2023), and as such, it is considered as an indicator of poor nutrient conditions (Navarrete et al. 2015). Given the significant differences in the microbial inoculations, it is possible that a combination of reduced access to rhizobia, fewer aggregate microbial benefits, as well as increased pathogen prevalence may have contributed to the worse plant performance in drought-treated soil.

Our root microbiome analysis did not find strong compositional and diversity differences due to drought-treated soil, likely because we only sequenced the microbiomes of live plants, which reduced our ability to detect microbes that may have contributed to differential mortality between climate-treated soils (e.g., survivorship bias). Furthermore, it is possible that there were more differences in the rhizosphere of plants early in the experiment, reflected in the strong decrease in performance of younger plants, but that these compositional differences were reduced over time by the mortality of affected plants and the selective environment of the growth chamber and the plants.

### Rhizobia inoculation reduced differences in plant performance due to climate-legacy

We added rhizobia to the soil to determine if mutualists could buffer plants against microbially-mediated negative effects of climate and found that the inoculation erased plant performance differences between climate-treated soils. If abundant mutualistic microbes reduce variation in plant performance and growth due to climate-driven differences in the microbiome, other organisms relying on those plants for nutrition or habitat will also experience less disruption, and this stabilizing effect could cascade across trophic levels (Kagata and Ohgushi 2006, Scherber et al. 2010). Thus, when partners are prevalent, mutualism can stabilize ecological dynamics against the indirect effects of climate change.

The relative abundance of compatible rhizobia in the un-inoculated field soil was very low (Boyle et al. 2024, Supplemental Figure 3), and in these conditions, climate-treated microbes altered plant performance. So, performance differences may have disappeared because of the increased abundance and accessibility of rhizobial mutualists to the host early in life, or potentially due to microbe-microbe interactions, both of which can affect host performance (Bakker et al. 2014, May and Nelson 2014, Burghardt et al. 2022). For example, a relatively high abundance of added rhizobia could have outcompeted detrimental microbes for resources or access to the host, or even altered the host as a niche via priority effects (Boyle et al. 2021, Debray et al. 2022, Burghardt and diCenzo 2023).

While performance was homogenized across climate treatments with the added rhizobia, plants with added rhizobia tended to do worse than plants with no added rhizobia in biotic soil, suggesting that the inoculated rhizobium strain was not a high-quality partner. This was not unexpected, as *S. meliloti* 1021 is well-characterized as a mediocre partner with *Medicago truncatula* (Terpolilli et al. 2008, Batstone et al. 2020a). Furthermore, the inserted green fluorescence protein (GFP) in our added strain of *Sinorhizobium meliloti* 1021-71 may make the microbe grow more slowly than non-modified strains (Rang et al. 2003) and may have required relatively more energy and resources from the plant. Another possibility is that our collected *M. lupulina* genotype is locally adapted to the rhizobia with which it co-occurs, and local rhizobia provided a greater benefit than the added strain (Batstone et al. 2020a). Our results indicate mutualists can reduce the effect of background variation in the microbiome if they are prevalent enough, however, for a truly protective effect they may need to match or supersede the quality of local mutualists.

### Mutualism quality and symbiotic patterns were robust to climate legacy

The maintenance of mutualism under changing environmental contexts will be important for interacting species’ fitnesses under future climate change, as well as continued ecosystem services (Kiers et al. 2010). Climate change is predicted to make mutualisms vulnerable to shifts towards antagonism, partner switches and losses, and mutualism abandonment (Brown 1997, Sachs and Simms 2006, Kiers et al. 2010). Our results demonstrated that heat- and drought-treated field rhizobia maintained their partner quality and ability to nodulate a legume host (Figure 3). The most immediate threat to legume-rhizobia symbioses may be the lower abundance or loss of a preferred partner, where the loss of a specialized mutualist means the other partner has few replacements and replacements are lower-quality (Froehlich et al. 2023).

In general, differences in the soil were not very predictive of differences in the root microbiomes (Supplemental Figures 3-4). Previous results found the most significant differences from climate treatments were due to drought and in the fungal community (Boyle et al. 2024), yet surprisingly, this did not manifest in significantly different fungal rhizospheres. One reason for this difference may be that many strongly affected fungi did not form a close symbiosis with *Medicago lupulina*, as most fungi in the soil were saprotrophs and most mycorrhizal fungi were ectomycorrhizal (Boyle et al. 2024). Moreover, it appears that there is increasingly strong host selection from the soil to the rhizosphere to the nodule; a pattern that has been observed in other legumes (Miranda Sánchez et al. 2016), suggestive of a deterministic microbiome (Lundberg et al. 2012). This pattern may arise due to factors such as host morphology, host rewards and byproducts, and host immune response, all of which can select for a subset of microbes from the broader microbial community (Zipfel and Oldroyd 2017, VanWallendael et al. 2022). Filtering mechanisms may increase plant performance if it also ensures recruitment of beneficial microbes and the exclusion of harmful microbes (Werner and Kiers 2015), but, in and across changing environments, the microbes to target and filter may be different (Caballol et al. 2024). An interesting next step would be to investigate the differences in how generalist or specialist legumes respond to microbially-mediated effects of climate, and the benefits and drawbacks of each strategy. While we observed a strongly deterministic root microbiome of living *M. lupulina* plants, it is also clear that when the ‘right’ microbes are affected by climate, legumes can still suffer lower plant performance and higher mortality.

## Conclusions

Plants experienced detrimental indirect effects of climate change through their microbiome, with the strongest effects early in life and due to the legacy of drought. Repeated drought events eroded the soil microbiome’s benefits under both drought stress and wet conditions, potentially reducing future plant resistance and resilience to drought. Indirect negative effects of drought on plant growth and fitness may challenge ecosystem productivity and function as well as agricultural yield. Additional access to rhizobia mutualists reduced microbially-mediated differences in plant performance due to climate. Our results motivate further inquiry into whether the addition of good-quality mutualists could buffer plants against this variation in the microbiome and potentially provide plants with increased stability under new climate regimes, with relevant applications in managed systems like agriculture and restoration sites. Our study highlights the complexity of plant response to components of climate change like extreme drought, and the threat of indirect microbially-mediated climate effects.

## Acknowledgements

We thank past and present KSR staff, Anna Simonsen for generating the seed line, Daniel Gage for providing the rhizobia strain, Chris Carlson for his help collecting soil and thoughtful discussions, Jessica Underwood for her help planting and applying treatments, and Bill Cole and Tom Gludovacz for growth chamber help. This work was funded by the Government of Canada through Genome Canada and Ontario Genomics (OGI-211), NSERC Discovery grants to IE, JRS, and MEF, the Canada Foundation for Innovation, and the University of Toronto’s Center for Global Change Science. We also acknowledge Genome Quebec for their sequencing facilities.

