## Supplemental for "Mutualism mediates legume response to microbial climate legacies"

Supplemental Information

Fertilizer recipe

Mix all ingredients in 500ml of distilled water, autoclave, wait for it to cool and then give each plant 1000ul. Ingredients are 2 ml MgSO_4_, 0.2 ml NaCl, 1.7 ml K_2_HPO_4_, 1.384 ml K_2_SO_4_, 2.4765 ml CaCl_2_, 0.235 ml Ca(NO_3_), 0.08 ml KNO_3_, 1.25 ml FeEDTA, 0.05 ml MnSO_4_, 0.05 ml CuSO_4_, 0.05 ml ZnSO_4_, 0.05 ml H_3_BO_3_, 0.05 ml Na_2_MoO_4_.

Experiment 2: Preparation and application of soil and rhizobia inoculations

We set aside 300 ml of fresh soil from each plot for biotic inoculation treatments, and another 300 ml for the abiotic soil treatments. We added 10 ml of live inoculation soil to the top of each conetainer, comprising 8% of the total volume of media available to plants. We added soil directly to maintain the integrity of fungal structures and diversity of microbes. We autoclaved the soil for abiotic treatments three times with a 30 minute sterilization length. Autoclaved soil became a smaller volume due to lost water content, and we used all the soil by adding 6 ml per plant. On the sixth day post-planting, we prepared an inoculation of *S. meliloti* 1021-71 in tryptone-yeast media at an optical density of 0.1, and applied 1 ml to plants with added rhizobia treatments. We added 1 ml of sterile tryptone-yeast media to plants without the rhizobia treatment to control for the presence of growth medium.

Experiment 2: Extracting the rhizosphere and nodule microbes

For sequenced plants, we first shook off the excess root media and separated aboveground biomass. We cut then the roots into 5 cm sections and placed them in a 15 ml falcon tube with 5 ml of cooled 1X phosphate-buffered saline (PBS) (plants with a lot of root mass were placed in 10 ml of PBS to allow full root submersion). We sonicated the roots for 15 minutes to dislodge the rhizosphere, then removed roots and centrifuged the PBS containing the rhizosphere and root-attached turface at 2500 g for 5 minutes to pellet microbes. The supernatant was discarded, and the pellet was kept at -20 °C until extraction. We counted and cut the nodules from the roots and surface-sterilized them with bleach and ethanol, and rinsed them with distilled water. We pooled nodules per plant and kept them at -20 °C until extraction. Then, we dried the plants’ above and below ground biomasses and weighed them.

We extracted the microbial DNA from the rhizosphere and the pooled nodules separately using QIAGEN DNeasy PowerSoil Pro Kits. The rhizosphere was sequenced for both ITS (primer pair ITS1FP2-58A2RP3) and 16S V4 region (primer pair 515F-806R) reads, while the pooled nodule samples (n = 67, since not all plants had nodules) were sequenced for the same 16S V4 region at Genome Quebec (Montréal, Canada) using Illumina MiSeq PE 250bp 16S rRNA amplicon sequencing. Additionally, we sequenced *S. meliloti* 1021-71 as a point of comparison.

Experiment 2: Microbiome processing

We used Quantitative Insights Into Microbial Ecology 2 (QIIME2) v.2022.2 (Bolyen et al. 2019). We trimmed the sequences for quality. For 16S reads we trimmed the left 20 bp from forward and reverse reads and truncated reads at 240 bp, while for ITS reads we trimmed the left 25 bp for forward sequences, trimmed 20 bp for reverse sequences, and truncated at 240 bp for both forward and reverse reads. We denoised the sequences with DADA2 (Callahan et al. 2016) into amplicon sequence variants (ASVs). We removed ASVs that had fewer than 10 reads across all samples, and assigned taxonomy using the ‘sklearn’ feature classifier (Pedregosa et al. 2011) with the 2021 Greengenes 16S V4 region reference for bacteria (McDonald et al. 2012), and UNITE version 9.0 with dynamic clustering of global and 97% singletons for fungi (Abarenkov et al. 2022). After assigning taxonomy, the bacterial rhizosphere had 8,058 ASVs with 3,055,989 reads and a median 26,923 reads/sample. The fungal rhizosphere had 1,016 ASVs and 2,902,562 total reads, and a median of 25,278 reads/sample. Nodule samples had 269 ASVs with 1,842,801 total reads, and median 28,518 reads/sample. Then we filtered out reads assigned as chloroplasts and mitochondria to remove plant DNA. After filtering, the bacterial rhizosphere retained 8,019 ASVs and 3,043,867 total reads, the fungal rhizosphere retained all ASVs and reads, and the nodule data retained 237 ASVs and 1,114,982 reads. We rarefied bacterial and fungal rhizosphere samples to 15,000 reads and 12,400 reads respectively, and rarefied nodule bacteria to 1000 reads. After rarefaction, we had 113 samples for the bacterial rhizosphere, 115 samples for the fungal rhizosphere, and 58 samples for nodules. Finally, we constructed phylogenies for bacterial reads using QIIME2’s MAFFT (Katoh & Standley 2013) and FastTree (Price et al. 2010) functions to obtain rooted trees.

Tables

**Supplemental Table 1. Effect of climate legacy on legume performance is context dependent.**

| **Condition dependency** | **Aboveground biomass (log g)** | | **Belowground biomass (log g)** | | **Final branch number** | | **Death** | |
| --- | --- | --- | --- | --- | --- | --- | --- | --- |
| *Predictors* | *Wald Chisq* | *p* | *Wald Chisq* | *p* | *Wald Chisq* | *p* | *Wald Chisq* | *p* |
| (Intercept) | 2971 | **<0.001** | 1401 | **<0.001** | 287 | **<0.001** | 0.00 | 1.00 |
| Historical soil condition | 20.0 | **<0.001** | 18.0 | **<0.001** | 4.64 | **0.031** | 0.604 | 0.437 |
| Experimental water condition | 18.5 | **<0.001** | 1.53 | 0.216 | 15.4 | **<0.001** | 9.58 | **0.002** |
| Historical: Experimental conditions | 5.98 | **0.015** | 7.44 | **0.006** | 2.10 | 0.148 | 0.162 | 0.687 |

**Supplemental Table 2. Effect of climate legacy on plant performance given biotic soil and no added rhizobia.**

| **Biotic, no added rhizobia** | **Leaf number over time (log)** | | **Aboveground biomass (log g)** | | **Belowground biomass (log g)** | | **Death** | |
| --- | --- | --- | --- | --- | --- | --- | --- | --- |
| *Predictors* | *Wald Chisq* | *p* | *Wald Chisq* | *p* | *Wald Chisq* | *p* | *Wald Chisq* | *p* |
| (Intercept) | 100 | **<0.001** | 606 | **<0.001** | 702 | **<0.001** | 13.7 | **<0.001** |
| Heat legacy | 0.00 | 0.945 | 0.052 | 0.820 | 0.260 | 0.610 | 1.72 | 0.190 |
| Drought legacy | 15.4 | **<0.001** | 1.34 | 0.248 | 1.93 | 0.165 | 4.33 | **0.037** |
| Heat: Drought legacies | 5.37 | **0.020** | 0.088 | 0.767 | 0.538 | 0.463 | 2.86 | 0.091 |
| Days after germination | 6699 | **<0.001** | NA | NA | NA | NA | NA | NA |
|  | **Nodule number (counts)** | | | **Nodule number (zeroes)** | | |  | |
| *Predictors* | *Estimate (std error)* | *z-value* | *p* | *Estimate (std error)* | *z-value* | *p* |  |  |
| (Intercept) | 2.93 (0.300) | 9.79 | **<0.001** | 0.546 (0.476) | 1.15 | 0.251 |  |  |
| Heat legacy [Yes] | -0.611 (0.435) | -1.41 | 0.159 | 0.723 (0.610) | 1.18 | 0.236 |  |  |
| Drought legacy [Yes] | -0.134 (0.418) | -0.320 | 0.749 | 0.521 (0.599) | 0.870 | 0.384 |  |  |
| Block [2] | -0.323 (0.089) | -3.64 | **<0.001** | -0.031 (0.422) | -0.072 | 0.942 |  |  |
| Block [3] | 0.278 (0.076) | 3.67 | **<0.001** | -0.639 (0.410) | -1.56 | 0.119 |  |  |
| Heat legacy [Yes]:Drought legacy [Yes] | 0.746 (0.604) | 1.23 | 0.217 | -0.740 (0.869) | -0.851 | 0.395 |  |  |

**Supplemental Table 3. Effect of climate legacy on plant performance and symbiosis given sterilized soil and no added rhizobia.**

| **Sterilized, no added rhizobia** | **Aboveground biomass (log g)** | | **Belowground biomass (log g)** | | **Death** | |
| --- | --- | --- | --- | --- | --- | --- |
| *Predictors* | *Wald Chisq* | *p* | *Wald Chisq* | *p* | *Wald Chisq* | *p* |
| (Intercept) | 569 | **<0.001** | 682 | **<0.001** | 11.6 | **<0.001** |
| Heat legacy | 0.00 | 0.992 | 0.231 | 0.631 | 0.653 | 0.419 |
| Drought legacy | 0.218 | 0.641 | 0.450 | 0.503 | 2.57 | 0.109 |
| Heat: Drought legacies | 1.56 | 0.212 | 1.13 | 0.287 | 0.186 | 0.666 |

**Supplemental Table 4. Effect of climate legacy on plant performance and symbiosis given biotic soil with added rhizobia.**

| **Biotic, added rhizobia** | **Leaf number over time (log)** | | **Aboveground biomass (log g)** | | **Belowground biomass (log g)** | | **Nodule number (log)** | | **Nodule presence** | | **Death** | |
| --- | --- | --- | --- | --- | --- | --- | --- | --- | --- | --- | --- | --- |
| *Predictors* | *Wald Chisq* | *p* | *Wald Chisq* | *p* | *Wald Chisq* | *p* | *Wald Chisq* | *p* | *Wald Chisq* | *p* | *Wald Chisq* | *p* |
| (Intercept) | 87.4 | **<0.001** | 409 | **<0.001** | 431 | **<0.001** | 22.8 | **<0.001** | 1.57 | 0.210 | 14.8 | **<0.001** |
| Heat legacy | 0.089 | 0.764 | 0.195 | 0.659 | 0.824 | 0.364 | 2.15 | 0.143 | 2.83 | 0.092 | 0.062 | 0.803 |
| Drought legacy | 1.47 | 0.225 | 0.077 | 0.782 | 0.169 | 0.681 | 0.071 | 0.791 | 1.15 | 0.284 | 0.00 | 0.994 |
| Heat: Drought legacies | 1.75 | 0.186 | 0.056 | 0.813 | 0.007 | 0.936 | 0.584 | 0.445 | 1.55 | 0.214 | 0.154 | 0.695 |
| Days after germination | 4375 | **<0.001** | NA | NA | NA | NA | NA | NA | NA | NA | NA | NA |

**Supplemental Table 5. Effect of climate legacy on plant performance and symbiosis given sterilized soil with added rhizobia.**

| **Sterilized, added rhizobia** | **Aboveground biomass (log g)** | | **Belowground biomass (log g)** | | **Nodule number (log)** | | **Nodule presence** | | **Death** | |
| --- | --- | --- | --- | --- | --- | --- | --- | --- | --- | --- |
| *Predictors* | *Wald Chisq* | *p* | *Wald Chisq* | *p* | *Wald Chisq* | *p* | *Wald Chisq* | *p* | *Wald Chisq* | *p* |
| (Intercept) | 460 | **<0.001** | 721 | **<0.001** | 29.0 | **<0.001** | 5.80 | **0.016** | 8.02 | **0.005** |
| Heat legacy | 0.593 | 0.441 | 1.53 | 0.217 | 1.34 | 0.246 | 1.09 | 0.295 | 0.008 | 0.927 |
| Drought legacy | 2.88 | 0.090 | 3.55 | 0.059 | 0.149 | 0.699 | 0.459 | 0.498 | 0.721 | 0.396 |
| Heat: Drought legacies | 1.73 | 0.188 | 3.27 | 0.071 | 0.802 | 0.371 | 0.161 | 0.688 | 0.270 | 0.603 |

**Supplemental Table 6. Effect of soil sterilization and added rhizobia on plant performance and symbiosis.**

| **Main treatments** | **Aboveground biomass (log g)** | | **Belowground biomass (log g)** | | **Death** | |
| --- | --- | --- | --- | --- | --- | --- |
| *Predictors* | *Wald Chisq* | *p* | *Wald Chisq* | *p* | *Wald Chisq* | *p* |
| (Intercept) | 1575 | **<0.001** | 1670 | **<0.001** | 32.8 | **<0.001** |
| Sterilization | 2.00 | 0.158 | 0.133 | 0.715 | 0.329 | 0.566 |
| Rhizobia | 6.66 | **0.010** | 5.23 | **0.022** | 0.431 | 0.511 |
| Sterilization:Rhizobia | 9.43 | **0.002** | 6.64 | **0.010** | 1.71 | 0.192 |
|  | **Nodule number (counts)** | | | **Nodule number (zeroes)** | | |
| *Predictors* | *Estimate (std error)* | *z-value* | *p* | *Estimate (std error)* | *z-value* | *p* |
| (Intercept) | 2.65 (0.079) | 33.6 | **<0.001** | -1.57 (0.356) | -4.41 | **<0.001** |
| Sterilization [Biotic] | -0.227 (0.036) | -6.40 | **<0.001** | 1.14 (0.373) | 3.04 | **0.002** |
| Rhizobia [None added] | 0.263 (0.038) | 6.86 | **<0.001** | 1.70 (0.247) | 6.87 | **<0.001** |
| Block [2] | 0.332 (0.041) | 8.03 | **<0.001** | -0.569 (0.278) | -2.04 | **0.041** |
| Block [3] | 0.388 (0.039) | 9.92 | **<0.001** | -1.02 (0.283) | -3.61 | **<0.001** |

**Supplemental Table 7. Bacterial rhizosphere alpha diversity metrics and community composition.**

| **Bacterial rhizosphere** | **Observed richness** | | **Shannon** | | **Evenness** | | **Community composition** | | | **Best and worst plants: community composition** | | |
| --- | --- | --- | --- | --- | --- | --- | --- | --- | --- | --- | --- | --- |
| *Predictors* | *Wald Chisq* | *p* | *Wald Chisq* | *p* | *Wald Chisq* | *p* | *F* | *R^2^* | *p* | *F* | *R^2^* | *p* |
| (Intercept) | 145 | **<0.001** | 748 | **<0.001** | 1560 | **<0.001** | NA | NA | NA | NA | NA | NA |
| Heat legacy | 4.08 | **0.043** | 2.59 | 0.107 | 1.05 | 0.305 | 1.01 | 0.009 | 0.411 | 1.29 | 0.027 | 0.079 |
| Drought legacy | 0.077 | 0.782 | 0.023 | 0.880 | 0.024 | 0.878 | 0.949 | 0.008 | 0.550 | 0.916 | 0.019 | 0.589 |
| Rhizobia | 1.68 | 0.195 | 2.17 | 0.141 | 1.87 | 0.171 | 1.35 | 0.012 | **0.047** | 1.09 | 0.023 | 0.318 |
| Heat: Drought legacies | 0.470 | 0.493 | 0.874 | 0.350 | 0.719 | 0.396 | 0.995 | 0.009 | 0.440 | 0.997 | 0.021 | 0.451 |
| Performance category | NA | NA | NA | NA | NA | NA | NA | NA | NA | 4.22 | 0.087 | **<0.001** |

**Supplemental Table 8. Fungal rhizosphere alpha diversity metrics and community composition.**

| **Fungal rhizosphere** | **Observed richness** | | **Shannon** | | **Evenness** | | **Community composition** | | | **Best and worst plants: community composition** | | |
| --- | --- | --- | --- | --- | --- | --- | --- | --- | --- | --- | --- | --- |
| *Predictors* | *Wald Chisq* | *p* | *Wald Chisq* | *p* | *F* | *R^2^* | *p* | *R^2^* | *p* | *F* | *R^2^* | *p* |
| (Intercept) | 77.5 | **<0.001** | 125 | **<0.001** | 150 | **<0.001** | NA | NA | NA | NA | NA | NA |
| Heat legacy | 2.91 | 0.088 | 0.648 | 0.421 | 0.011 | 0.916 | 0.774 | 0.007 | 0.642 | 1.23 | 0.03 | 0.181 |
| Drought legacy | 1.78 | 0.182 | 0.00 | 0.996 | 0.180 | 0.672 | 0.524 | 0.005 | 0.928 | 1.08 | 0.02 | 0.328 |
| Rhizobia | 0.031 | 0.861 | 0.299 | 0.584 | 0.955 | 0.328 | 0.622 | 0.006 | 0.834 | 1.00 | 0.02 | 0.415 |
| Heat: Drought legacies | 0.130 | 0.718 | 0.085 | 0.771 | 0.015 | 0.904 | 1.13 | 0.010 | 0.272 | 0.97 | 0.02 | 0.493 |
| Performance category | NA | NA | NA | NA | NA | NA | NA | NA | NA | 1.22 | 0.03 | 0.20 |

**Supplemental Table 9. *Sinorhizobium* strain composition and alpha diversity.**

| ***Sinorhizobium* in nodules** | **Observed richness** | | **Shannon (binomial)** | | **Evenness** | | **Community composition** | | |
| --- | --- | --- | --- | --- | --- | --- | --- | --- | --- |
| *Predictors* | *Wald Chisq* | *p* | *Wald Chisq* | *p* | *Wald Chisq* | *p* | *F* | *R^2^* | *p* |
| (Intercept) | 0.861 | 0.354 | 2.47 | 0.116 | 6.11 | **0.013** | NA | NA | NA |
| Heat legacy | 0.295 | 0.587 | 0.897 | 0.344 | 0.185 | 0.667 | 1.28 | 0.023 | 0.232 |
| Drought legacy | 0.438 | 0.508 | 2.41 | 0.121 | 0.052 | 0.820 | 0.746 | 0.014 | 0.505 |
| Nodule number | 1.30 | 0.255 | 4.89 | **0.027** | 0.104 | 0.748 | 0.515 | 0.009 | 0.676 |
| Heat: Drought legacies | 0.130 | 0.718 | 0.781 | 0.377 | 0.083 | 0.774 | 0.393 | 0.007 | 0.822 |
| Rhizobia | NA | NA | NA | NA | NA | NA | 0.588 | 0.011 | 0.648 |

Figures

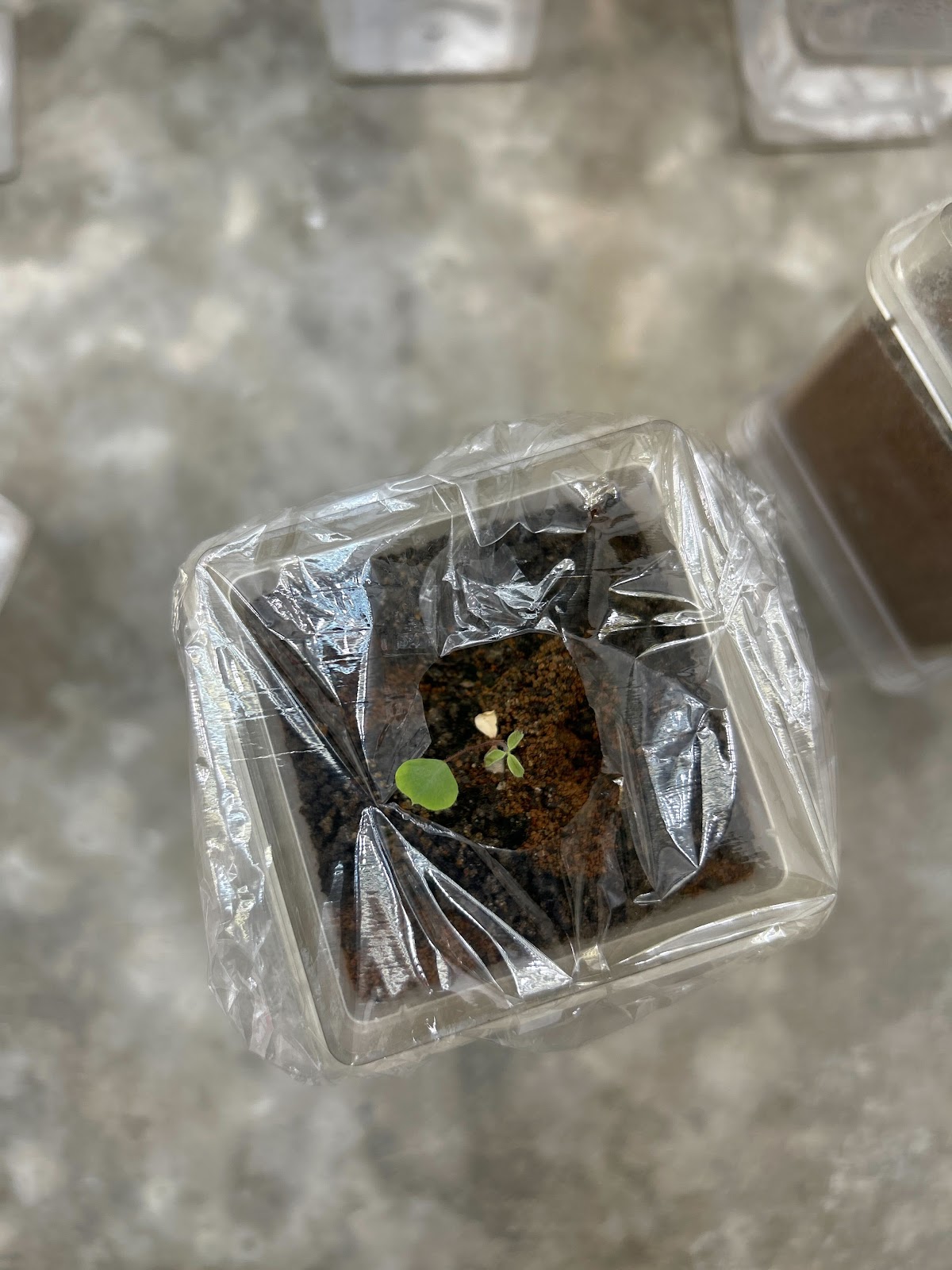

**Supplemental Figure 1. Plastic wrap cover on a well-watered plant, with a hole to accommodate plant growth.**

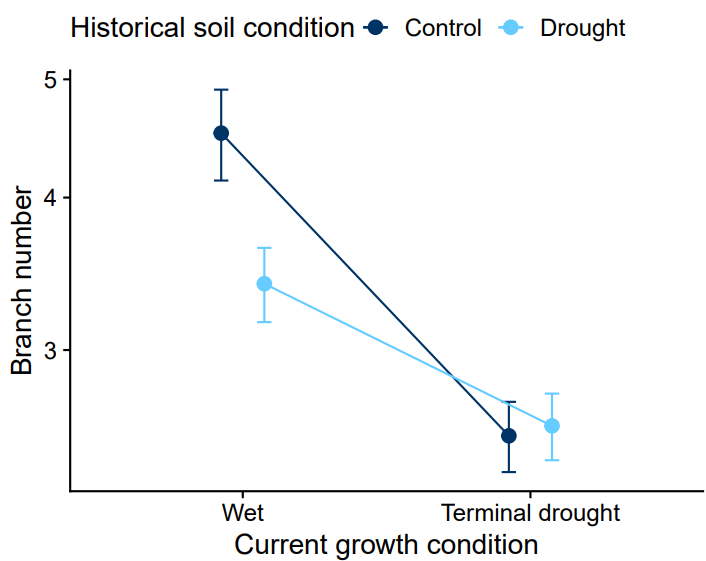

**Supplemental Figure 2. Branch number decreased due to terminal drought and drought-treated microbes.** The y-axis is scaled with a log_10_ transformation.

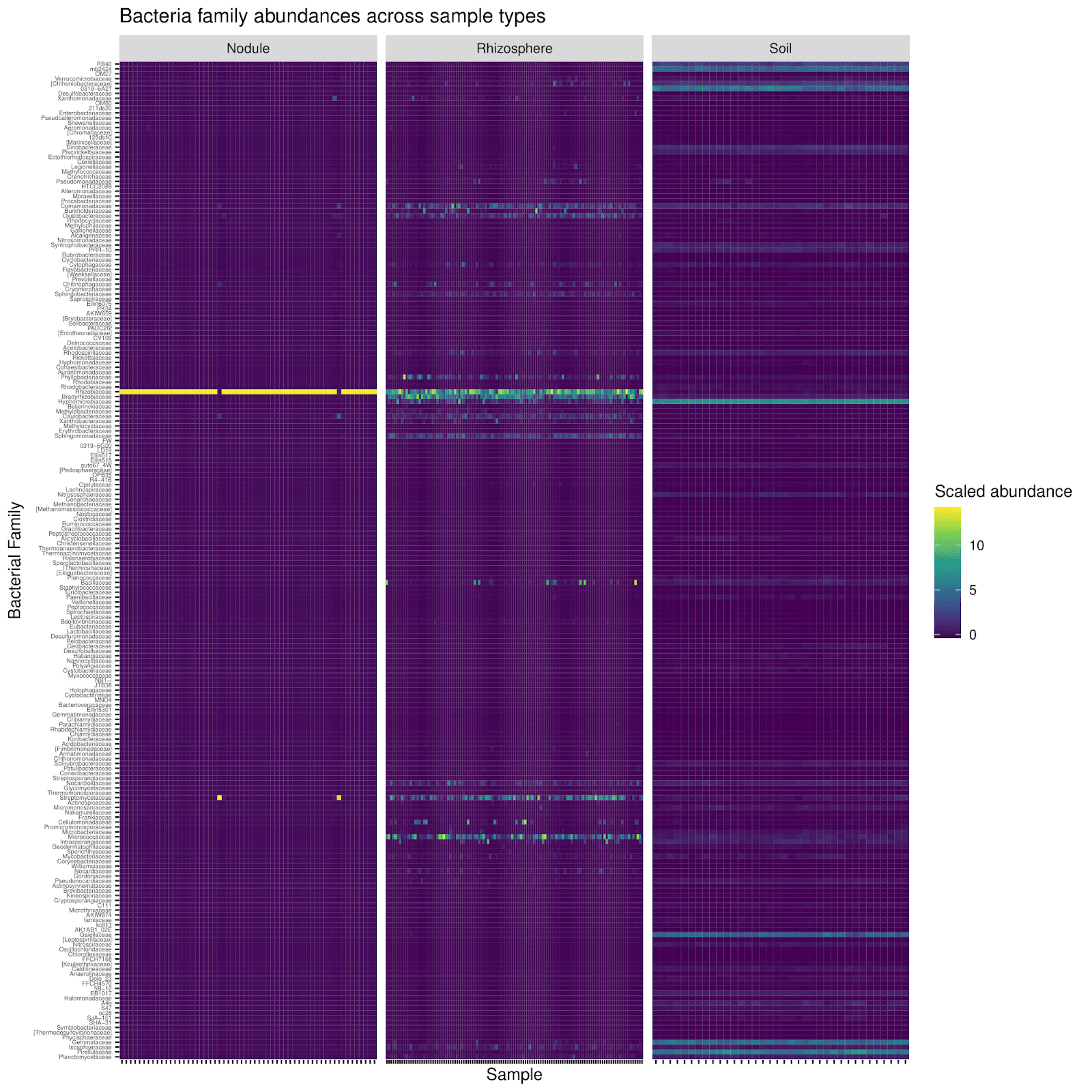

**Supplemental Figure 3. Scaled abundance of bacterial families across sample types.** Sample types include all samples with >1000 reads from the nodules and rhizospheres of sequenced plants, and the inoculation soil (showing the sequenced three subsamples per array plot). Bacterial families are ordered by their phylogenetic proximity. Abundances were scaled within each sample type.

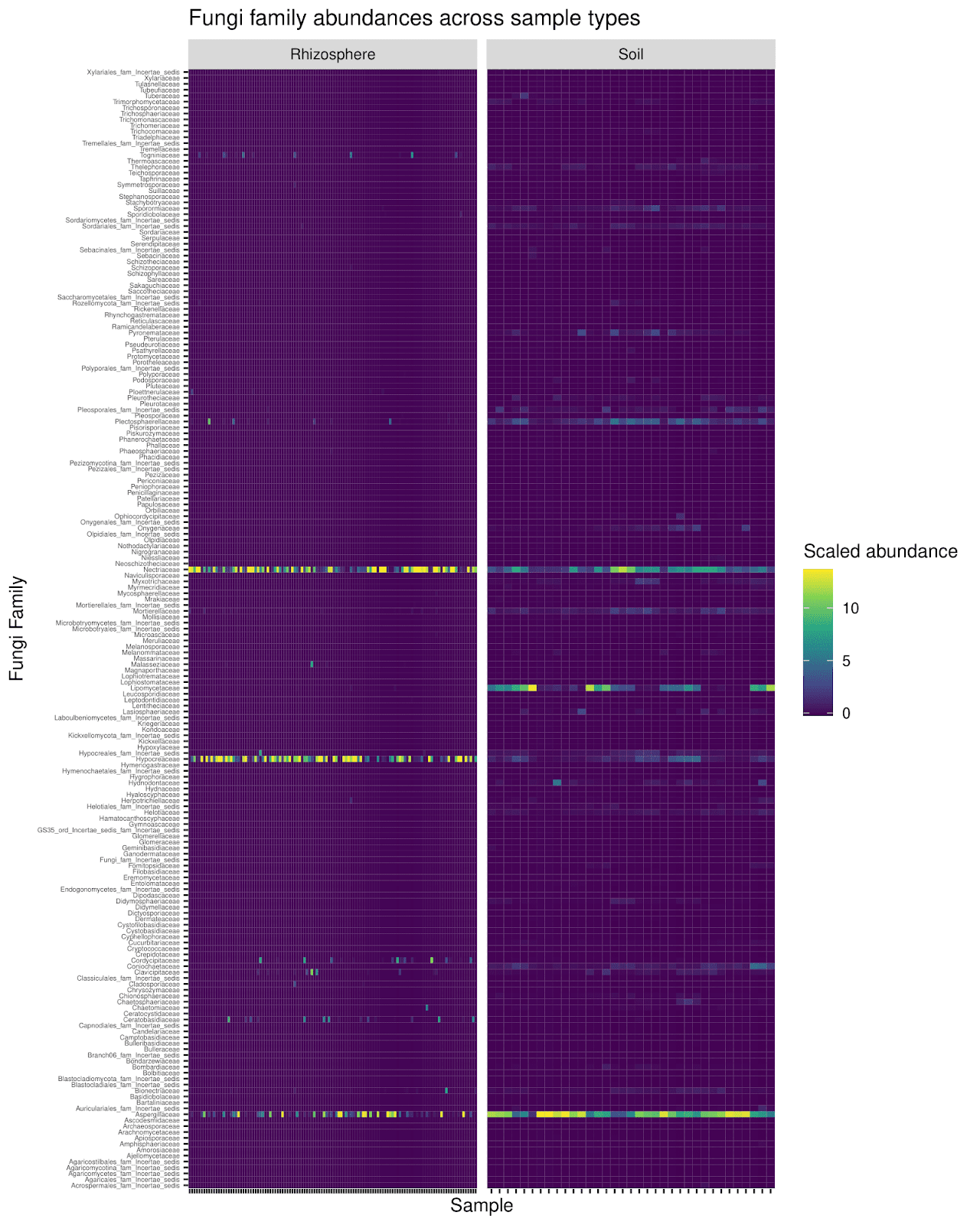

**Supplemental Figure 4. Scaled abundance of fungal families across sample types.** Sample types include all samples with >1000 reads from rhizospheres of sequenced plants and the inoculation soil (showing the sequenced three subsamples per array plot). Abundances were scaled within each sample type.

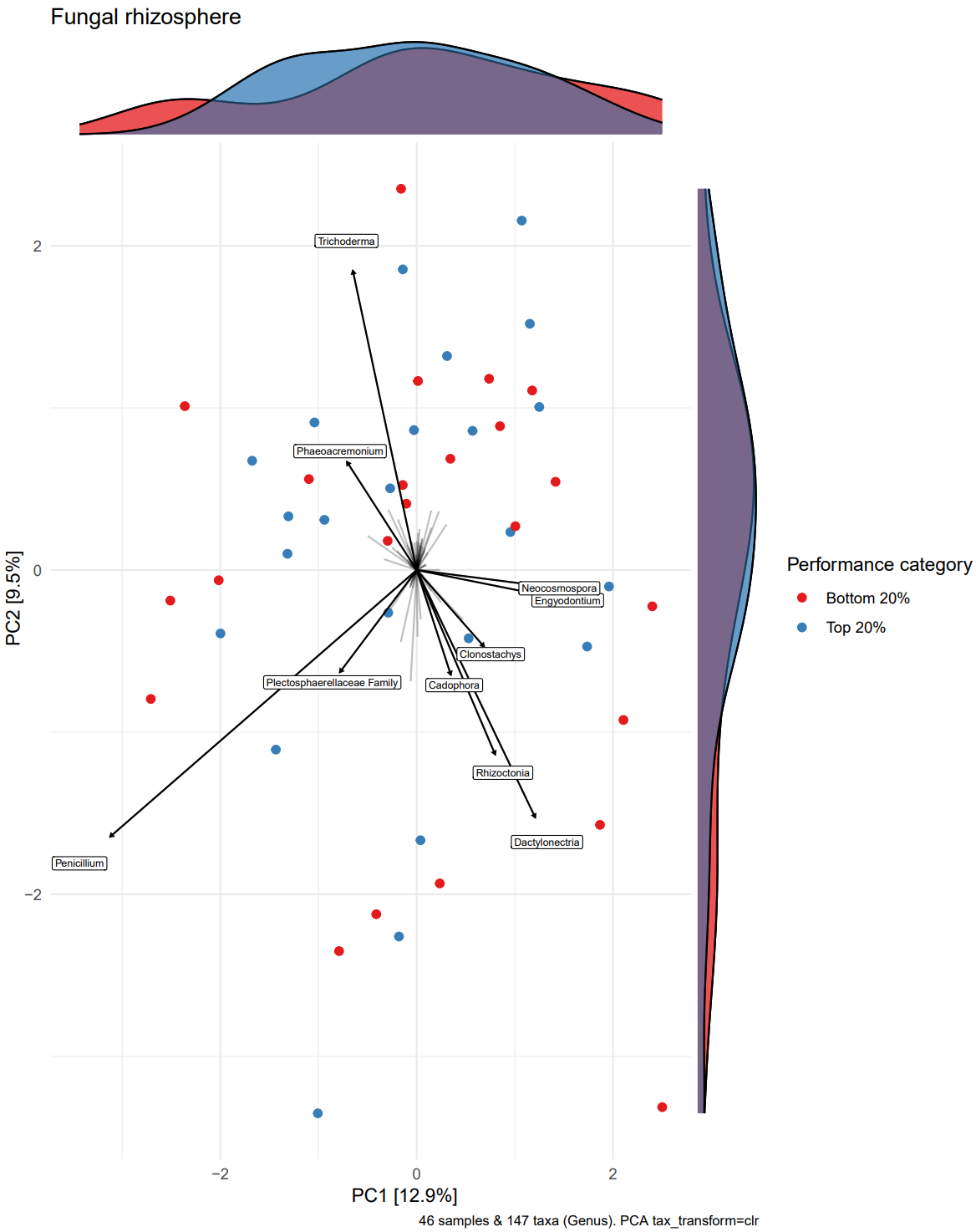

**Supplemental Figure 5. Microbes were not clearly correlated to performance differences in the fungal rhizosphere.** Here we use a PCA to show the center log ratio transformed fungal rhizosphere composition of the top 20% best performing and bottom 20% worst performing plants. Performance was quantified by aboveground biomass (g).
